# Regulation of interstitial fluid flow along adventitia of vasculature by heartbeat and respiration

**DOI:** 10.1101/2022.10.18.512678

**Authors:** Hongyi Li, Bei Li, Wenqi Luo, Xi Qi, You Hao, Chaozhi Yang, Wenqing Li, Jiazheng Li, Zhen Hua, Tan Guo, Zhijian Zheng, Xue Yu, Lei Liu, Jianping Zhao, Tiantian Li, Dahai Huang, Jun Hu, Zongmin Li, Fang Wang, Hua Li, Chao Ma, Fusui Ji

## Abstract

Converging studies showed interstitial fluid (ISF) adjacent to blood vessels flows along adventitia of vasculature into heart and lungs. We aim to reveal circulatory pathways and regulatory mechanism of such adventitial ISF flow in rat model. By MRI, real-time fluorescent imaging, micro-CT and histological analysis, ISF was found to flow in adventitial matrix surrounded by fascia and along systemic vessels into heart, then flow into lungs via pulmonary arteries and back to heart via pulmonary veins, which was neither perivascular tissues nor blood or lymphatic vessels. Under physiological conditions, speckle-like adventitial ISF flow rate was positively correlated with heart rate, increased when holding breath, became pulsative during heavy breathing. During cardiac or respiratory cycle, each dilation or contraction of heart or lungs can generate to-and-fro adventitial ISF flow along femoral veins. Discovered regulatory mechanisms of adventitial ISF flow along vasculature by heart and lungs will revolutionize understanding of cardiovascular system.

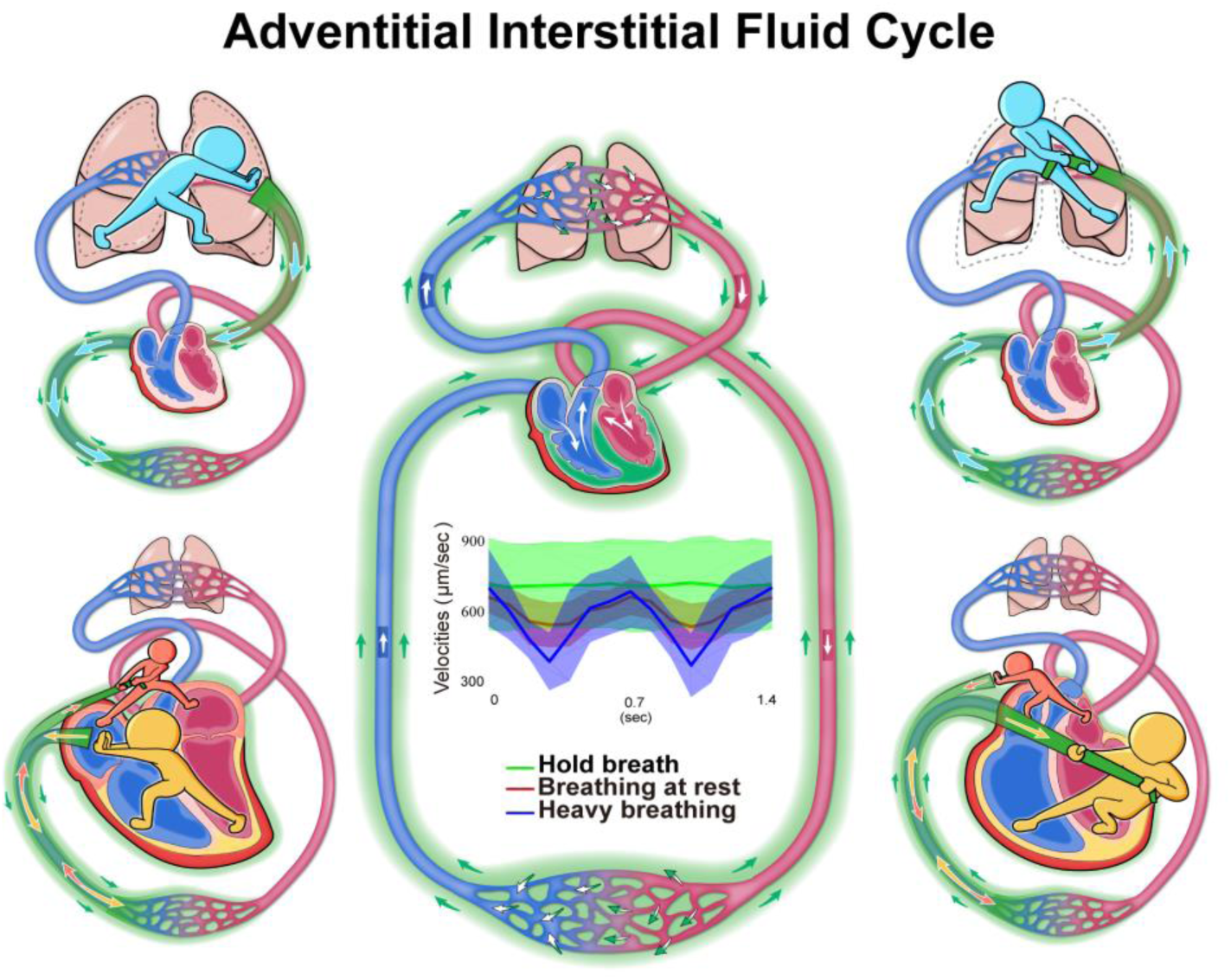
Graphic abstract of adventitial interstitial fluid cycle by heartbeat and respiration. The heart and lungs can work together to regulate the ISF (green arrow) adjacent to blood vessels to flow centripetally in an adventitial pathway along the major arteries and veins of the systemic vasculature to the epicardium and myocardium, then flow into the lungs via the pulmonary arteries and back to the heart via the pulmonary veins, forming an adventitial ISF circulatory network. Under physiological conditions, the speckle-like adventitial ISF flow rate was positively correlated with heart rate, increased when holding breath, and became pulsative during heavy breathing. During respiratory cycle, each expansion of the lungs during inflation generates a centrifugal flow of adventitial ISF along the systemic veins, while each contraction of the lungs during deflation generates a centripetal adventitial ISF flow. During cardiac cycle, each dilation of atria or ventricles generates a centripetal flow of adventitial ISF along the systemic veins, while each contraction of atria or ventricles generates a centrifugal adventitial ISF flow.

## Introduction

Since *Willian Harvey* discovered blood circulation in 1628, and the *Starling’s hypothesis* was proposed in 1896, the motion of extracellular fluid throughout the body is believed to include a rapid circulation of blood and lymph, and a constant movement of the ISF through extracellular matrix after filtrated from the capillary walls. To date, long-standing efforts have found that the ISF could not only diffuse around tissue cells but also flow along blood vessels or in the interstitial compartment of certain tissues or organs over a long distance, such as the perivascular spaces (PVS) in the brain, thymus, liver and several other organs ^1–9^, the interstitial tissue channels between capillaries and initial lymphatic vessels ^10,11^, the interstitial spaces of tumors ^12,13^, etc. These ISF flows are distributed regionally near the arterioles, capillaries and venules at the ends of the vascular tree, and regulated by hydrostatic and osmotic pressure differences ^14^, capillary filtration coefficient, pumping by lymphatic system ^15^, shear stresses generated by blood flow in the vascular microenvironment like the hypothesis of perivascular pump ^16,17^, and etc. Recently, converging studies have shown that the ISF adjacent to arteries and veins can also flow along the adventitia of the vasculature into the heart and lungs under an unknown regulatory mechanism.

In brain by intracisternal injection of ovalbumin-conjugated fluorescent tracer, the ISF flow was found in the PVS along the penetrating arteries, arterioles, capillaries and venules as well as the large-caliber draining veins of mice ^6,8^. In the body by interstitial injection of water-soluble low-molecular-weight fluorescent or paramagnetic tracers into perivascular tissues, our research group found that the ISF was illustrated by the tracers to flow longitudinally in the adventitial connective tissues along major arteries and veins from the extremities to the heart and lungs in animal models and human subjects ^18–21^. In rabbits by the injection of the fluorescent tracer into the ankle hypodermis of rabbits, such adventitial ISF was visualized to flow along the saphenous and femoral veins, the inferior vena cava in the abdomen and thorax, and into the heart, while diffusing into the perivascular tissues along the way, and the capillaries, venules and lymphatic vessels within them and back into blood circulation ^18^. The calculated velocity of the continuous fluorescent adventitial ISF flow along the femoral veins was 3.6∼15.6 mm/s ^21^. In the pulmonary circulation of rabbits, the adventitial ISF was found to flow along the pulmonary vein from the lungs toward the heart ^21,22^. The arterial adventitial flow was demonstrated in the femoral artery of the rabbits, mice and the posterior tibial artery of humans ^18,19,21,23^. In the human cadaver experiments, the ISF from the thumb hypodermis could be “pulled” via the intact adventitial pathways along the superficial and deep veins of arm, axillary sheath and superior vena cava into the atrial walls when the heart was repeatedly compressed by an automatic cardiac compressor ^20^. Given arborescent vasculature exists everywhere throughout the body, elucidating the ISF flow along blood vessels will provide a novel mechanism for regulating the fluid environment in diverse tissues or organs by means of vascular vessels in humans and animals.

Using improved tracer imaging and quantitative speckle tracking techniques, we studied the whereabouts of the ISF flow along the adventitia of vascular vessels, the directions of the adventitial ISF flow along the blood vessels in the systemic and pulmonary circulation, and the mechanisms by which the heart and lungs regulate adventitial ISF flow in the rat.

## Results

### The adventitial pathways originating from the right saphenous vessels by MRI, real-time fluorescence stereomicroscopy, or SMCT and their differences from angiography

Consistent with our previous studies ^18,21,23,24^, after administration of the water-soluble tracers into the perivascular tissues or adventitia of the large-caliber vascular vessels of lower limbs, it was found that the tracers would centripetally flow and visualize the arteries and veins starting from the administration point to the heart. When compared to angiography by intraluminal injection, it was the adventitial tissues distributed along the vessels were visualized by the adventitial infused tracers, rather than the blood within the lumen.

By MRI at 5min after the hypodermic injection of the Gd-DTPA into the right ankle dermis, it was clearly found that the contrast enhanced the arterial and venous vessels of the right lower limb, but not the arteries and veins of the contralateral lower limb (Fig. S1B, Video 1). In comparison, the intravascular contrast had visualized the arteries and veins of both right and left lower limbs at 5min after the tail vein injection (Fig. S1F, Video 2). At 10-20min after the injection, the adventitial contrast gradually enhanced the arteries and veins of the contralateral lower limb, the aorta and IVC of the abdominal and thoracic cavities, kidney, and bladder, indicating that the adventitial contrast flowed along the vascular vessels and meanwhile entered the blood circulation (Fig. S1C, S1D). At 40-50min after the adventitial infusion or intravenous injection, the visualization of blood vessels began to weaken (Fig. S1D, S1H). The results of dynamic MRI scanning suggested that the adventitial infused tracer enhanced the adventitial tissues along the blood vessels during about 5min after the administration, and then gradually entered the blood circulation.

By real-time fluorescence stereomicroscopy or SMCT at 3min after the adventitial infusion, either the adventitial FluoNa or silver nitrate from the saphenous vessels was found to flow centripetally along the saphenous vessels (Fig. 1B, 1K, 1G, 1N) and stained the proximal end (Fig. 1C, 1L) but not the distal end (Fig. 1D, 1M) of the vessels. By analyzing the real-time frozen sections, the fluorescence intensity of the proximal site was significantly stronger than those of the distal section in the adventitial infusion group (Fig. 1C, 1D, 1E). By comparison, there were no significant differences found between the proximal and distal ends in the angiography group (Fig. 1H, 1I, 1J). The SMCT results showed that the intravenous cavity of the saphenous vein was stained by the intravenously injected silver nitrate (Fig. 1N, 1O, 1P).

**Figure 1.**
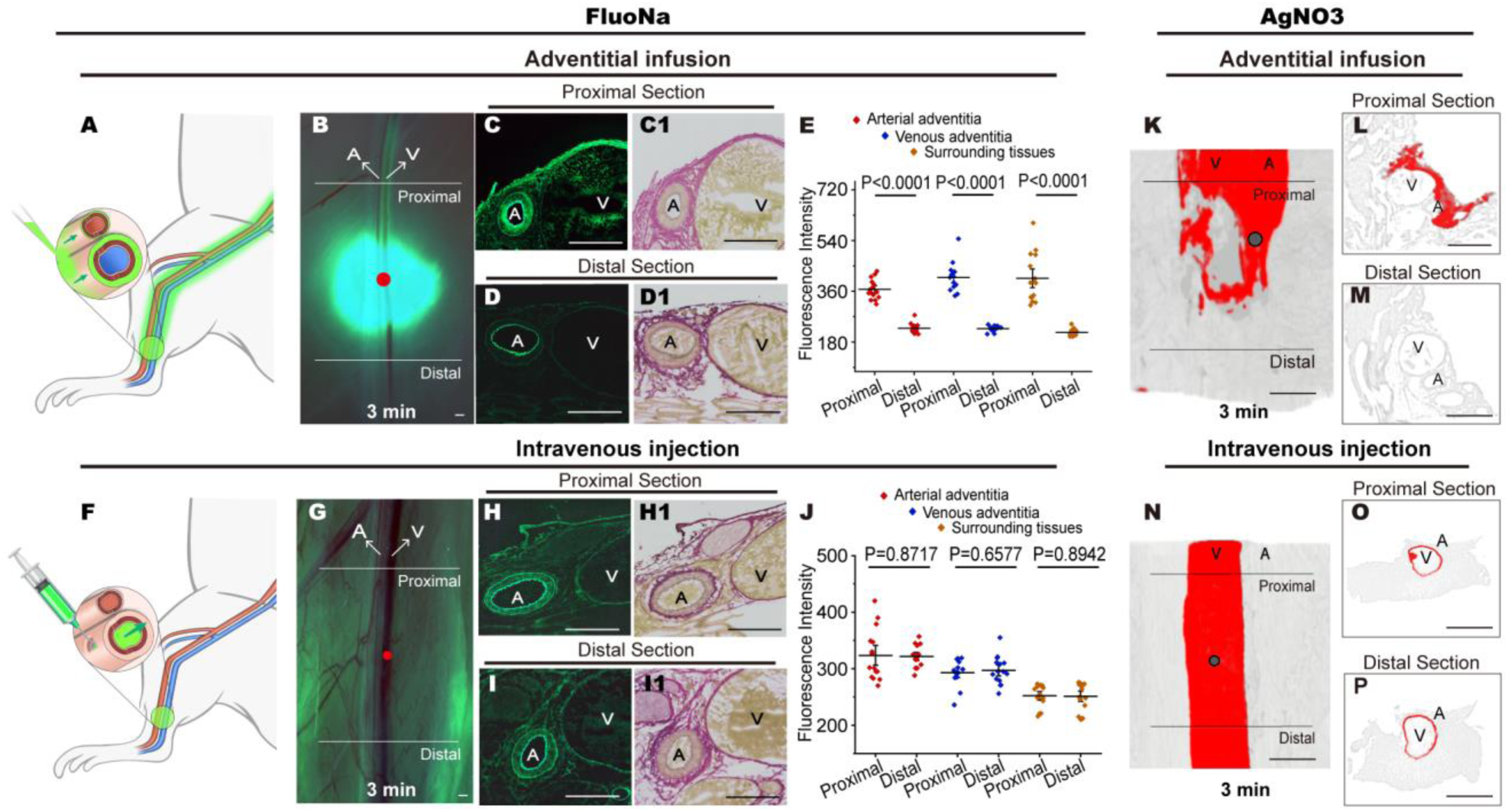
Comparison of adventitial infusion and intravenous injection of the FluoNa and Silver nitrate in the lower limbs. By fluorescence stereomicroscopy, **A**, **B** The FluoNa was infused into the adventitial pathways of the saphenous vessels, diffused around the infusion site (red point) and flowed centripetally along the vessels. Cross-sectional images showed the proximal (**C, C1**) but not the distal (**D, D1**) end of the adventitial pathways was stained by the FluoNa. The analysis of fluorescence intensity showed that the fluorescent signals of the arterial and venous adventitia, and the tissues surrounding the vessels in the proximal end were significantly stronger than those of distal end (**E**). **F**, **G** The FluoNa was intravenously injected into the saphenous vein (red point). There were no significant differences of fluorescence intensity between the proximal (**H, H1**) and distal (**I, I1**) end of the adventitial pathways (**J**). Using SMCT, the adventitially infused silver nitrate was found to flow along the proximal but not distal end of the adventitial pathways (**K**, **L**, **M**). The intravenous silver nitrate enhanced the proximal and distal end of the vein together (**N**, **O**, **P**). Scale bar: 500μm. The solid lines in (**B**, **G**, **K**, **N**) indicated the sites of cross section at the proximal or distal end. **A**, artery. **V**, vein. In (**E**, **J**), *t*-test, mean ± SEM, n = 3 rats.

### Quantifying the longitudinal adventitial ISF flow along femoral artery and vein by fluorescent tracer

To identify whether the location of adventitia ISF flow is in the perivascular tissues or adventitial matrix, we surgically striped of the perivascular tissues along a segment of femoral vessels and used real-time fluorescence stereomicroscopy to study the flow of the adventitial infused FluoNa (4µL, 0.1 g/L) in the adventitia of the isolated artery and vein. At 30sec after the adventitial infusion into the distal adventitial pathways on the saphenous vessels, the cross-sectional views showed that the FluoNa had stained the adventitia surrounding the isolated femoral artery and vein (Fig. S2B, S2B1).

When the adventitia along the femoral vein and artery were disrupted by type I collagenase, it was found that the distal FluoNa cannot flow through and stain these femoral adventitial channels as well as the arterial wall (Figs. 2D, 2D1, S2C, S2C1). At the same time, the angiography confirmed that both the femoral artery and vein were unobstructed (Fig. 2E, Videos 3, 4). Histological analysis of successive sections showed that the femoral artery and vein did not contain the vasa vasorum running continuously along the long axis of the vessels (Fig. S3A-J). The lymphatic vessels were also present in the perivascular connective tissues but not within the adventitia of blood vessels (Fig. S3K, S3K1, S3K2, S3L). Thus, the tunica adventitia surrounded by fascia, rather than the conduit-like vasorum and lymphatics, were the histological structures for the longitudinal adventitial ISF flow. Estimated by micro-CT, the numbers of the adventitial fibers under its surrounding fascia were calculated to be around 2504 (range from 2315 to 2796) along the femoral vein, and 2605 (range from 2450 to 2750) along the femoral artery (Fig. 2Q).

**Figure 2.**
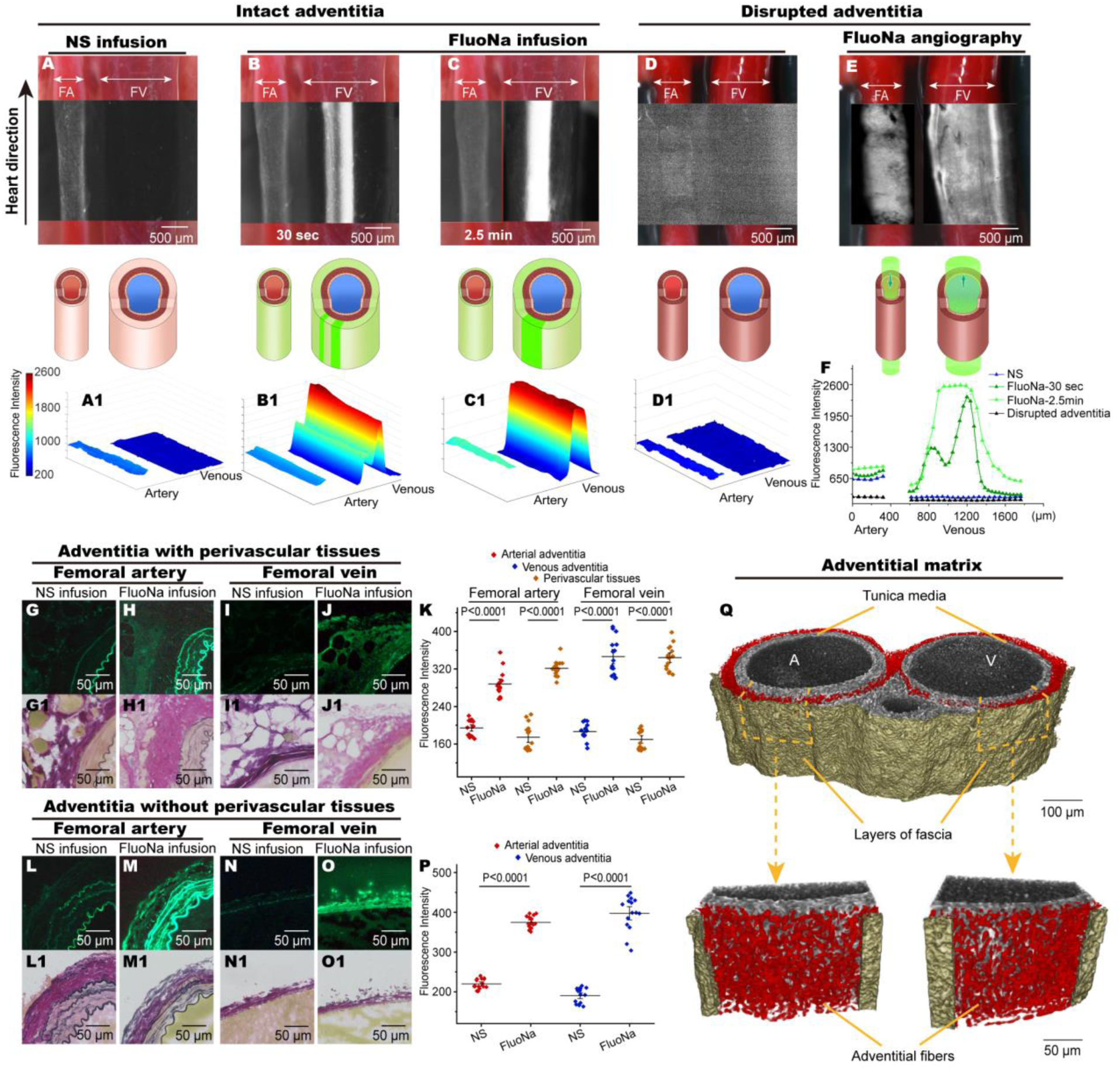
Quantifying the longitudinal adventitial ISF flow along the femoral artery and vein that were stripped of perivascular tissues. To clearly observe the longitudinal adventitial ISF flow, the femoral arteries and veins (**A**, **B**, **C**, **D**, **E**) were stripped of the perivascular connective tissues. Recorded by real-time fluorescence stereomicroscope, **A**, **A1** showed that the fluorescence intensity of a femoral artery and vein was weaker by the adventitial infusion of NS. After the adventitial infusion of FluoNa into the distal adventitial pathways of the saphenous vessels, two linear fluorescent lines in the adventitial pathway were found along the side of the femoral vein at 30sec (**B**) and converged into one wider fluorescent line channel at 2.5min (**C**). When the adventitia of femoral vessels was disrupted by collagenase I (**D**), the distal FluoNa cannot flow through and stain the adventitial walls of the artery and vein. Meanwhile, the contralateral venous angiography confirmed that this femoral artery (**D**) was unobstructed, and the ipsilateral venous angiography showed this femoral vein (**D**) was also unobstructed (**E**). **A1**, **B1**, **C1**, **D1** showed the intensity and distributions of the fluorescein along the arteries and veins detected by fluorescence stereomicroscopy. **F** showed that the changes in fluorescence intensity of the (**A1**, **B1**, **C1**, **D1**). **G**-**J** were the cross-sectional views of femoral vessels by fluorescence microscopy and showed the results of the fluorescent adventitial ISF in the adventitia of femoral vessels having stained the perivascular tissues as well. (**G1**-**J1) were the bright field corresponding to (G-J). L**-**O** showed the results of the fluorescent ISF flowing in the adventitia of femoral vessels without the perivascular tissues. (**L1**-**O1) were the bright field corresponding to (L-O). H, H1** showed that the arterial adventitia, connective tissues surrounding the artery, and even the media were all stained by the FluoNa from the distal, and the fluorescence intensity was significantly stronger (**K**) than that of the control group **G**. **J, J1** showed that the venous adventitia, connective tissues surrounding the vein, and even the media were all stained by the FluoNa from the distal, and the fluorescence intensity was significantly stronger (**K**) than that of the control group **I**. **M, M1** showed that the arterial adventitia and media can be stained by the distal FluoNa even in the absence of perivascular tissues, and the fluorescence intensity was significantly stronger (**P**) than that of the control group **L**. **O, O1** showed that the venous adventitia and media can be stained by the distal FluoNa in the absence of perivascular tissues, and the fluorescence intensity was significantly stronger (**P**) than that of the control group **N**. **Q** The reconstructed three-dimensional views of the femoral artery and vein. The numbers of the adventitial fibers between the fascia and tunica media were calculated to be around 2504 (range from 2315 to 2796) along the femoral vein, and 2605 (range from 2450 to 2750) along the femoral artery. **FA**, femoral artery. **FV**, femoral vein. The arrows marking FA or FV refer to the inner diameter of the FA or FV under bright field.

Following administration of the FluoNa by adventitial infusion onto the adventitia of distal vessels of the limbs under real-time stereomicroscopy with a high-resolution camera, two types of adventitial ISF flow were observed along the femoral vessels, early rapid flow, and subsequent speckle-like flow. Within 20 seconds after the start of the adventitial infusion, it was found that the FluoNa flowed rapidly along the femoral veins and formed two fluorescent lines along one side of the femoral vein. At subsequent 20-30 seconds when more FluoNa entered the adventitial pathways, the fluorescence intensity of these two fluorescent lines gradually increased (Fig. 2B, 2B1), changed from thin to wide, and merged into a wider fluorescent line at about 120-150 second (Figs. 2C, 2C1, S3A-D). Inside the wider fluorescent line along the femoral vein, the flow of the FluoNa was relatively stable and exhibited speckle-like between consecutive frames (Video 5). Limited by the current *in vivo* imaging technique, the longitudinal flow of fluorescent ISF in the femoral arterial adventitia was observed, but the speckle-like flow cannot be detected (Figs. 2C, 2C1, S3A).

In the relatively fixed HR and certain respiratory parameters, the flow rate measured in the adventitial pathways along the femoral veins was consistent when 1μl, 4μl and 16μl of infusion fluid of the FluoNa were administrated into the distal end of the adventitial pathways (Table S1). Then one of infusion doses, 4μl was selected for per administration in the studies.

Under physiological conditions with the HR of 290-460 bpm and certain respiratory parameters (RR: 90bpm, I/E: 1:2, TV: 4.0mL), the rapid flow of the FluoNa along the femoral vein was recorded within 20sec after the administration and the calculated flow rate was 1.3-5.8mm/sec (Fig. S4). At 2.5min after administration, the continuous speckle-like flow was dynamically recorded along the femoral veins and the velocity was measured by STV (Fig. S5A) ^25–28^. In the 10-minute observation, we typically tracked around 130,000 speckles (Table S2). The median velocity of the calculated speckles was around 588.2μm/sec (range 258.0 to 967.1μm/sec), and the mean velocity was 598.1±107.7μm/sec (Table S2). The velocity profile of the ISF flow field was uniform from the left edge to the right, from the bottom to the top (Fig. S5B-E), but not a parabola ^29–31^. By comparison with fluorescence angiography, the calculated velocity of the arterial blood flow was 188.5±57.6mm/sec, and the venous blood flow was 6.6±0.6mm/sec (n = 5 rats for each group). In contrast, the velocities of the ISF flow in tumor tissues or connective tissues were 0.1-6.3μm/sec ^32–34^.

The measured arterial and venous pressure were 98-112/80-88mmHg, 5-9mmHg under the HR of 360-430bpm (Table S3). By comparison, we found that only the FluoNa can exhibit the speckle-like flow along the adventitial pathways, while Rhodamine B and Indocyanine green cannot.

### Regulation of adventitial ISF flow along femoral vein by HR and respiration

Using STV under physiological conditions, the flow rate of the adventitial ISF along the femoral vein was found to be associated with the HR and respiration. When the HR increased from 150-450 bpm, the flow rate of the adventitial ISF also increased (Fig. 3B). When the apnea was present and the HR was 410-420 bpm, the median flow rate of the adventitial ISF was significantly faster than that of resting breathing (Fig. 3C, 3C1). When breathing heavily with large amount of TV (6.0mL), a pulsatile flow of the adventitial ISF was displayed, the frequency of which was the same as the heavy breath (Fig. 3C, 3C1).

**Figure 3.**
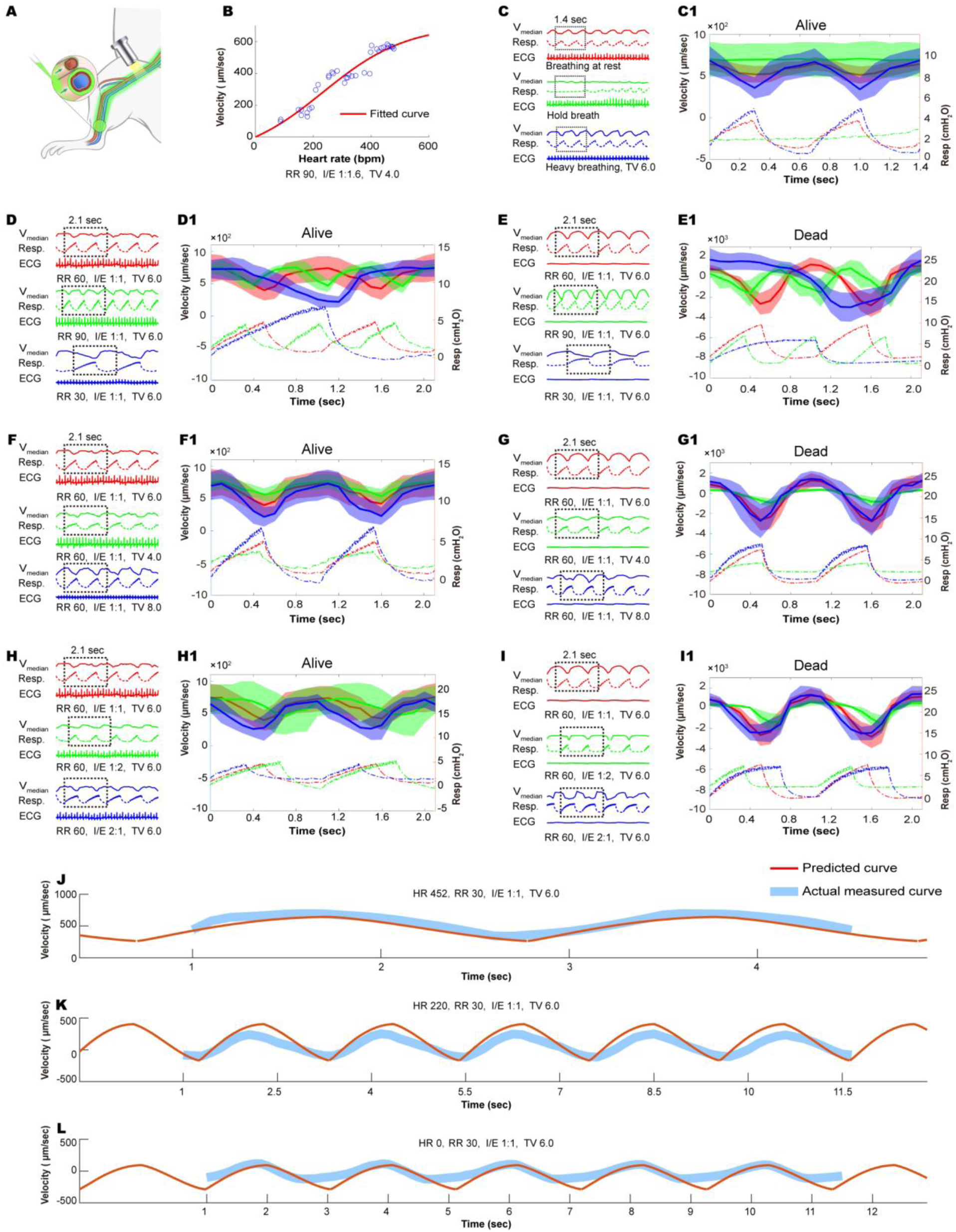
Regulation of adventitial ISF flow in the venous adventitial pathways by HR and respiration. **A** Illustration of the observation on the adventitial ISF flow in the venous adventitial pathways of the femoral vessels. **B** When the HR increased from 150-450 bpm, the flow rate of the adventitial ISF also increased (RR 90bpm, I/E 1:1, TV 4.0mL, n=1). The fitted curve was calculated by the cyclical dynamic equation. **C**, **C1** When the breath held, like apnea, the adventitial ISF flow rate increased by comparison with that at resting breath. A heavy breath induced a pulsatile flow at the same frequency as the heavy breath. **D**, **D1**, **E**, **E1** The frequency of the pulsed flow was consistent with the RR in alive or dead rats. **F**, **F1**, **G**, **G1** The greater the tidal volume, the greater the pulse amplitude of adventitial ISF flow in alive or dead rats. **H**, **H1**, **I**, **I1** The inspiration and expiration (I/E) ratio (1:1, 2:1, 1:2) determined the descending and ascending branches of the pulse in alive or dead rats. In alive rats, the inflation can decelerate the centripetal adventitial ISF flow. In dead rats, the inflation can cause a centrifugal adventitial ISF flow, while the deflation induced a centripetal adventitial ISF flow. Each group of (**C**, **D**, **E**, **F**, **G**, **H**, **I**) was sampled from 6 rats. The solid lines were the mean value and the shaded is the standard deviation of the median velocity of a total 6 rats in each group of **C1**, **D1**, **E1**, **F1**, **G1**, **H1**, **I1**, respectively. The dotted lines (Resp.) are the measured pressure on the surface of the body of a rat by the breathing band sensor, representing the changes of breathing. (**J**) was from Video 6, (**K**) was Video 7, and (**L**) was Video 8. The HR of (**J**) was around 452bpm, (**K**) was 220bpm, and (**L**) was zero. The ventilation parameters of (**J**), (**K**), and (**L**) were all the same, the RR was 30 breaths/min, I/E was 1:1, and tidal volume was 6.0mL. In either (**J**) or (**K**) or (**L**), the predicted curve (orange) calculated by this cyclical dynamic equation matched the actual measured curves (blue) of the adventitial ISF flow rate very well.

When the HR of rats was 390-420bpm, the frequency of the pulsed flow rate of the adventitial ISF was consistent with the RR (Fig. 3D, 3D1); at different tidal volumes, the greater the tidal volume is, the greater the pulse amplitude of the adventitial ISF flow rate (Fig. 3F, 3F1) is; I/E (1:1, 2:1, 1:2) determined the descending and ascending branches of the pulse (Fig. 3H, 3H1).

When the heart of rats stopped beating and the lungs were ventilated repeatedly, the act of inflation can cause a centrifugal flow of the adventitial ISF, while the act of deflation induced a centripetal flow of the adventitial ISF. Correspondingly, the effects of the RR, TV, and the I/E ratio were consistent with those under physiological conditions (Fig. 3E, 3E1, 3G, 3G1, 3I, 3I1). Meanwhile, the measured venous pressures were significantly reduced to 0-1mmHg by comparison with 5-7mmHg under physiological conditions (Table S3).

### The empirical dynamic equation for the regulation of adventitial ISF flow by heartbeat and respiration (Supplementary Note 1)

Assuming that the adventitial pathway with diverse types of interior channels can be simplified as a cylindrical channel, and the ISF is an incompressible and Newtonian fluid with a low Reynolds coefficient, the general form of the correlation between the velocity (*V*_*max*_) and the pressure gradients (Δ*p*) over a given length of the channel will be

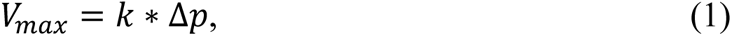

*k* is a continuous and determined by the properties of both fluid and adventitial pathways, such as the fluid’s viscosity, length of the pathways, the radius of the point (at different positions in the channel) to the center of the channel (the equation (6) in Supplementary Note 1).

According to our experimental findings, there are two cyclical driving forces regulating the adventitial ISF flow along the femoral veins, the heartbeat and respiration. Δ*p_heart_* represents the driving forces generated by the regularly cyclical deformations of the heart and is related with the frequency θ of the heartbeats. Δ*p_Lung_* represents the driving forces generated by the irregularly cyclical deformations of the lungs and is related with the respirations, including the respiratory frequency *F*, tidal volume *V*, inspiration-expiration ratio *R*. Therefore, we can get the empirical dynamic equation as

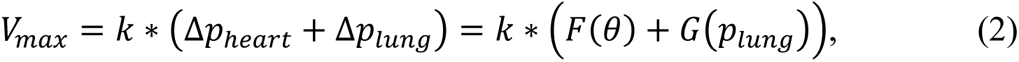

where F and G are the functions related to the heartbeats and respirations respectively. We can determine a function model depending on the observation and then derive the parameters of the function model via fitting method. The derivation process of the empirical equation was listed in the Supplementary Note 1. The final equation of *V*_*max*_ will be

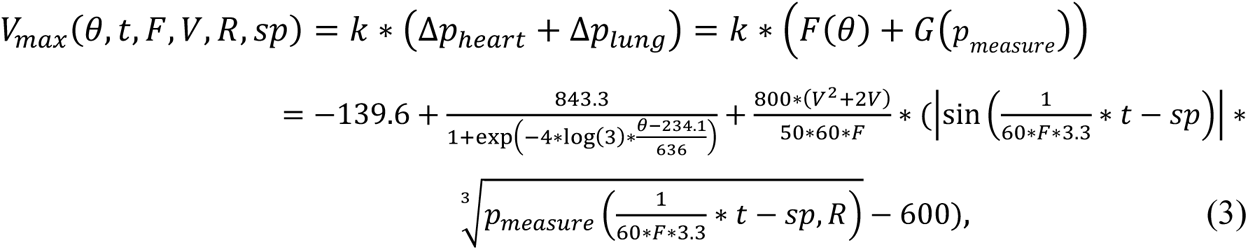

The parameter *sp* is referred to the starting phase (*sp*) of the irregular cycle of the respiration and is determined individually by each case. *t* is referred to time. By comparison, the predicted curve (orange) calculated by this cyclical dynamic equation matched the actual measured curves (blue) of the adventitial ISF flow rate very well (Fig. 3J-L, Videos 6-8).

Therefore, these revealed correlations between the adventitial ISF flow and the cyclical motions of heartbeat and respiration indicated that the ISF is able to pass through the interior channels (inter-channel) of the adventitial pathways in response to external forces. Considering the *k* represented the properties of a simplified cylindrical channel, we named the transport ability of the ISF flow through the diverse types of inter-channels of an adventitial pathway as hydraulic interfacial/inter-channel transportability (HIT, represented by *T*). The empirical dynamic equation (2) can be reformed as

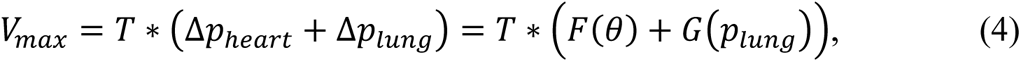

*T* is determined not only by *k* but also by the properties of the inter-channels of the adventitial pathways for fluid flow, such as the interfacial properties of the inter-channels of the adventitial pathways, the liquid content in the gel matrix, the connectivity of the fibrous scaffold of the adventitial matrix, temperature, etc. The HIT of diverse adventitial pathways needs further studies to be determined.

### To-and-fro motions of adventitial ISF along femoral vein during cardiac and respiratory cycle

By analyzing the changes of the mean velocity by STV, the effects of periodic deformations during the cardiac or respiratory cycle on the adventitial ISF flow along the femoral veins were observed. Under the actions of repeated chest compressions at intervals of 2-3sec in the freshly dead rats, the contraction and dilation of the heart were found to cause the same pulsatile flow of the adventitial ISF (Fig. 4A), while the venous pressure was 1-2mmHg (Table S3). When the chest cavity opened and the atria or ventricles were compressed at an interval of 10-15sec, it was found that every compression of the atria (Fig. 4B, 4B1) or ventricles (Fig. 4C, 4C1) induced a centrifugal motion of the adventitial ISF, and every dilation of the atria (Fig. 4B, 4B2) or ventricles (Fig. 4C, 4C2) caused a centripetal motion, while the venous pressure was 1-2mmHg (Table S3). In one respiratory cycle, the expansion of the lungs during inspiration caused the adventitial ISF to move centrifugally (Fig. 4D, 4D1), while the contraction of the lungs during expiration caused a centripetal movement of the adventitial ISF (Fig. 4D, 4D2). The venous pressure was measured to be 0-1mmHg under such one-shot ventilation (Table S3).

**Figure 4.**
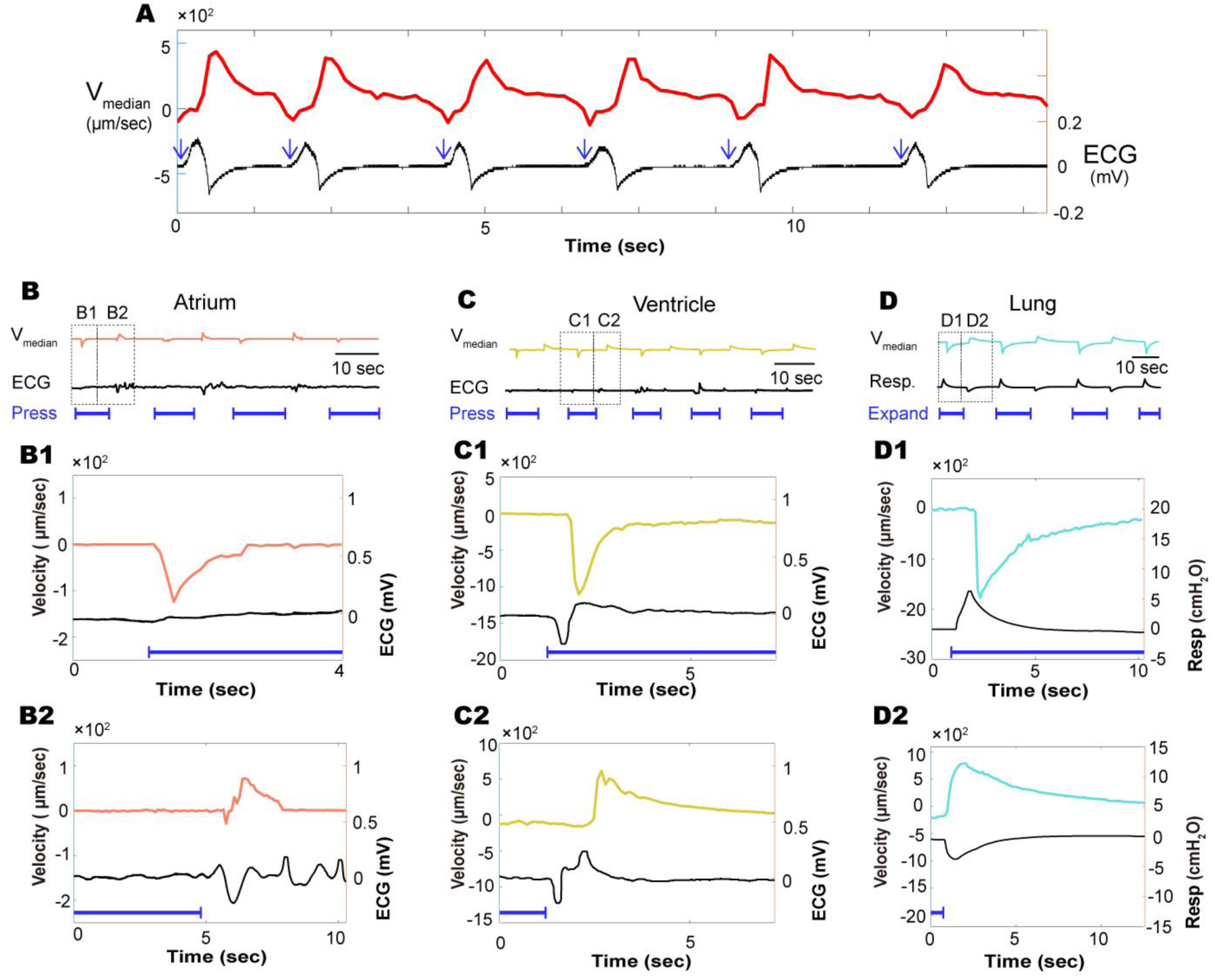
To-and-fro motion of adventitial ISF in the femoral adventitial pathways during a cardiac and respiratory cycle. **A** By repeated chest compressions at intervals of 2-3sec in the freshly dead rats, the contraction and dilation of the heart caused the same pulsatile flow of the adventitial ISF along the femoral veins. Blue arrow indicates the one-shot act of chest compression. When the chest cavity was opened, the atria or ventricles were continuously compressed for 10-15sec (blue line), and then released to allow the atria or ventricles to dilate on their own (**B**, **C**). **B1**, **C1** The compression of the atria or ventricles induced a centrifugal adventitial ISF flow. **B2**, **C2** The dilation of the atria or ventricles caused a centripetal adventitial ISF flow. **D** The lungs were expanded by the 5ml syringe for 10-15sec (blue line), and then contracted by the 5ml syringe for another 10-15sec. **D1** The expansion of the lungs induced a centrifugal adventitial ISF flow. **D2** The contraction of the lungs caused a centripetal adventitial ISF flow.

In the dead rats with open ventricular chambers, no blood flow from the apex was observed during the experiments, whether by repeated cardiac compressions or mechanical ventilations of lungs. Meanwhile, the measured arterial and venous blood pressures were 10.3±2.3/4.9±0.5mmHg and 0.7±0.1mmHg by cardiac compressions, or 3.8±1.0/3.7±1.0mmHg and 0.6±0.3mmHg by ventilations, respectively (Fig. S6). However, under repeated cardiac compressions, it was observed that the distal FluoNa was able to flow centripetally along the left femoral vessels but could not pass through the adventitial pathways of the right femoral vessels, which were disrupted by collagenase I (Fig. 5B, 5B1, 5E, 5E1). In contrast, under repeated ventilation of the lungs, the proximal FluoNa was observed to flow centrifugally along the left femoral vessels, but not through the adventitial pathways of the right femoral vessels, which had been disrupted by collagenase I as well (Fig. 5G, 5G1, 5J, 5J1). Moreover, even if the cardiac compressions or ventilations of the lungs could cause the mechanical fluctuations in the vascular walls of the lower limbs, the FluoNa in the adventitial pathways can flow centripetally driven by the heart or centrifugally driven by the lungs, respectively (Fig. 5A, 5D, 5F, 5I).

**Figure 5.**
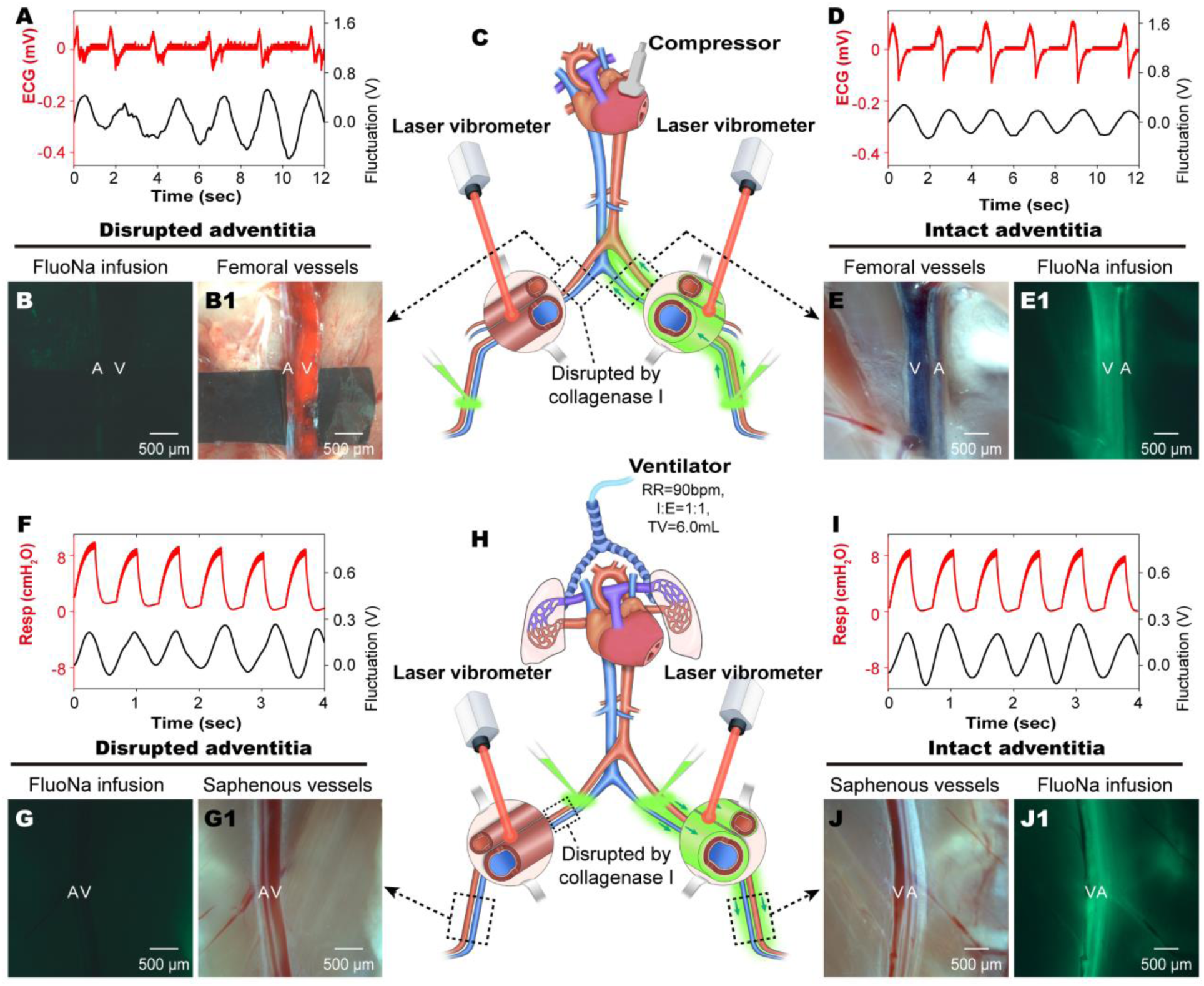
Illustration of the effects of the repeated cardiac compressions or ventilations on the adventitial flow in freshly dead rats with opened ventricular chambers. In the freshly dead rats, the apex of the heart was cut open to air so that the repeated cardiac compressions or ventilations cannot induce blood flow, and the mechanical motions of the vessel wall was detected by a laser vibrometer. The adventitial pathways along the right femoral vessels were disrupted by collagenase I, while the left remained intact (**C**, **H**). After the FluoNa was infused into the adventitial pathways on distal saphenous vessels of both sides, the heart was compressed repeatedly at 30-60bpm for 30 minutes in 6 rats while the lungs were not ventilated (**C**). **H** showed that the lungs were ventilated (RR 90bpm, I/E 1:1, TV 6.0mL) for 30 minutes in another 6 rats while the heart was not compressed after the FluoNa was infused into the adventitial pathways on the proximal femoral vessels of both sides. **B**, **B1** The distal FluoNa cannot flow centripetally through the right disrupted adventitial pathways. **E**, **E1** The distal FluoNa flowed centripetally and stained the intact adventitial pathways along the left femoral vessels. **G**, **G1** The proximal FluoNa cannot flow centrifugally through the right disrupted adventitial pathways. **J**, **J1** The proximal FluoNa flowed centrifugally and stained the intact adventitial pathways along the left saphenous vessels. **A**, **D**, **F**, **I** The mechanical motions were detected on the wall of both the right (**A**) and the left (**D**) femoral vessels. **A**, artery. **V**, vein.

When the root of the main pulmonary artery was clamped, the expansion and contraction of the lung can still cause the to-and-fro motions of the adventitial ISF along the femoral vein. However, after clamping of the thoracic segment of the inferior vena cava, the movements of the adventitial ISF along the femoral vein disappeared. These results suggested that effects of the lungs’ expansions and contractions on the adventitial ISF flow along the femoral venous vessels were achieved through the pulmonary venous vessels, rather than the pulmonary arteries.

### Qualitative observations on adventitial ISF flow pathways along the systemic and pulmonary arteries and veins by FluoNa

For the systemic vasculature, the adventitia-infused FluoNa from the right lower limb was found to stain the adventitial connective tissues of inferior vena cava (Fig. S7E-G), aorta (Fig. S7B-D), and the heart muscles of the left ventricle (Fig. 6B). The FluoNa infused into the adventitial pathways of the right axillary artery and vein (Fig. S7K-M), and the left carotid artery (Fig. S7H-J) was found to have stained the right anterior vena cava (Fig. S7N-P), aorta, and the heart. These data suggested that the adventitial ISF flow in the adventitial pathways centripetally along both arteries and veins of the systemic vasculature into the heart.

**Figure 6.**
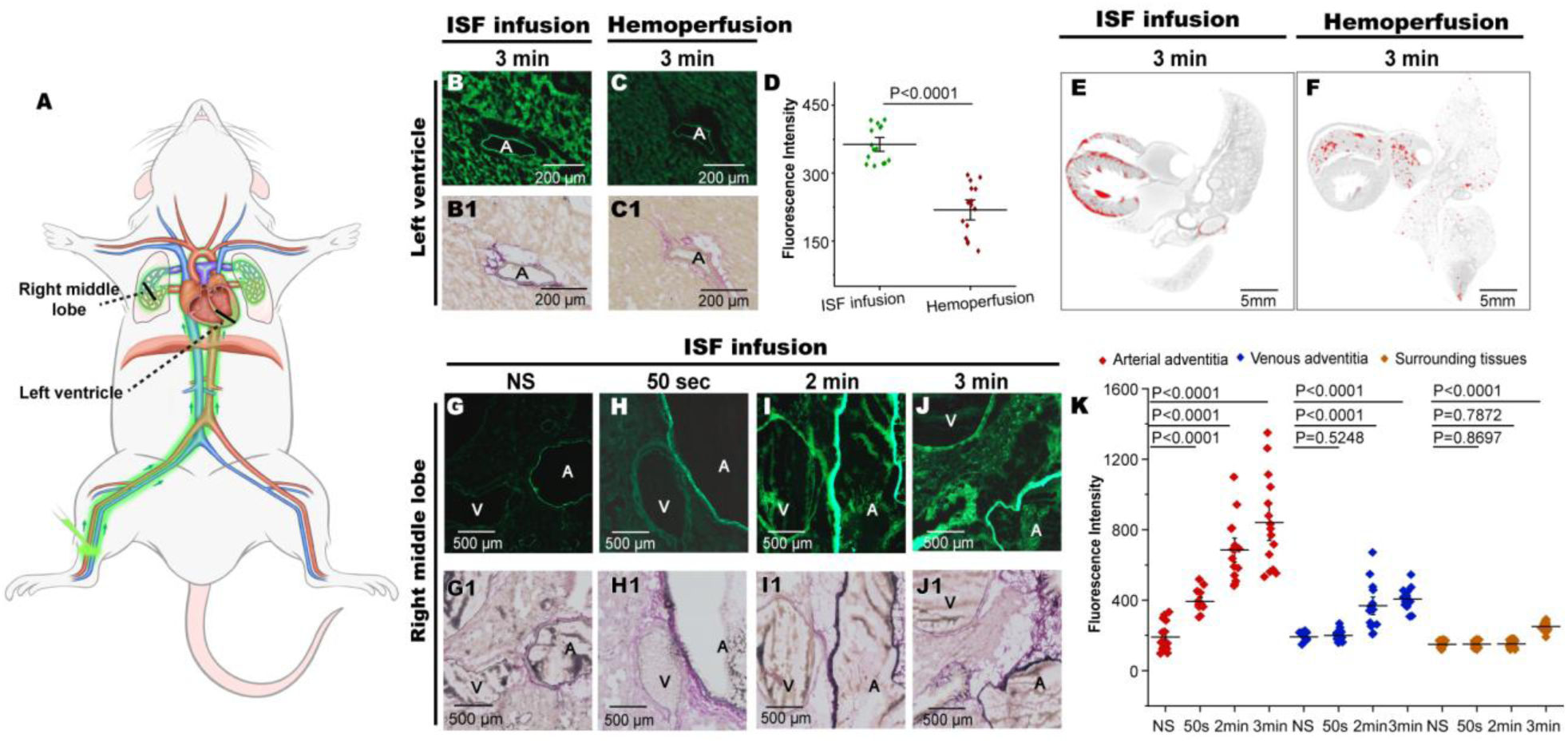
Qualitative observations on adventitial ISF perfusion and intravenous perfusion of the FluoNa or silver nitrate into the heart, and the circulatory cycle of adventitial ISF along the pulmonary vasculature. At 3min after the adventitial infusion of FluoNa from the adventitial pathways of the lower limb or the intravenous injection of FluoNa, the fluorescence intensity (**B**) of the myocardial tissues near the left ventricular epicardium by adventitial infusion (**A**) was significantly stronger than those by intravenous injection (**C**). (**B1**-**C1) were the bright field corresponding to (B-C).** By SMCT, the silver nitrate (red) from the adventitial pathways of the lower limb was found in the left and right ventricular muscles near the left and right ventricular epicardium (**E**) significantly but few in the lungs. In contrast, the silver nitrate by intravenous injection was found inside the right ventricle, some coronary arteries, and pulmonary vessels (**F**). To delineate the adventitial ISF cycle of the pulmonary vasculature, FluoNa was infused into the adventitial pathways of the lower limbs and detected in the middle lobe of the right lung at different times. **G, G1** The cross-sections showed a pulmonary artery and vein in the middle lobe and sampled at 50sec after the adventitial infusion of NS. **H, H1** At 50sec after the adventitial infusion, the distal FluoNa stained the adventitial pathway of the pulmonary artery but not the adventitial pathways of the vein and the perivascular tissues. **I, I1** At 2min after the adventitial infusion, the distal FluoNa stained the adventitial pathway of the pulmonary artery and vein but not the perivascular tissues. **J, J1** At 3min after the adventitial infusion, the distal FluoNa stained the adventitial pathway of the pulmonary artery and vein and the perivascular tissues. **K** The changes of the fluorescence intensity of the adventitial pathways in arteries, veins and the surrounding tissues, respectively. The findings indicated that the adventitial ISF flow cycle along the pulmonary vasculature is from the heart to the lungs along the pulmonary arteries, and from the lungs to the heart along the pulmonary veins. In (**D**, **K**), *t*-test, mean ± SEM, n = 3 rats. **A**, artery. **V**, vein. The sites of cross section in in heart (**B**, **C**) and lung (**G**-**J**) were pointed at (**A**) as the black solid lines.

For the pulmonary vasculature, it was also found that the adventitial FluoNa could move along the pulmonary artery toward the lungs (Fig. S8A, S8A1) or along the pulmonary vein toward the heart (Fig. S8B, S8B1). The findings indicated that the adventitial ISF flow cycle along the pulmonary vasculature was from the heart into the lungs via the pulmonary arteries and back to the heart via the pulmonary veins.

For the coronary arteries of the heart, the FluoNa, that was infused into the superficial tissues on the ventricular apex, was found to flow in the adventitial pathways along the coronary arteries toward the base of the heart (Fig. S10A-C, Video 9), suggesting that the upper part of the heart above the coronary sulcus might be the driver for the adventitial ISF flow along the coronary arteries of the ventricles.

The distributions in the myocardium of the adventitial infused tracers from the lower limbs were studies by the FluoNa or silver nitrate, respectively. At 3min after the administration of the FluoNa into the adventitia of saphenous vessels in lower limb, it was found that the fluorescence intensity in the myocardium (Fig. 6B, 6B1) by the adventitial infusion of the FluoNa was stronger than that of hemoperfusion by intravenous injection at 3min (Fig. 6C, 6D). The SMCT showed that the silver nitrate from the adventitial pathways along the saphenous vessels have entered the ventricular muscles (Fig. 6E, Video 10), the amount of which was much more than those by hemoperfusion (Fig. 6F). In addition, the silver nitrate from the adventitial pathways was mainly distributed in the subepicardial tissues. However, how to identify the adventitial ISF from the systemic vasculature circulate throughout the heart still needs more research.

### Qualitative observations on the adventitial ISF flow starting from the ends of the systemic vasculature, and the circulatory pathways of adventitial ISF from pulmonary arteries to veins by FluoNa

Does the adventitial ISF flow only under pathological conditions like extremity edema, or is it an inherent flow under physiological conditions? Using conventional angiography, we found that even without administration of the FluoNa directly into the adventitial pathways, the FluoNa from arterial blood still stained the adventitial pathways along both the femoral vein and artery after capillary exchange in the distal lower limb (Video 11). The results suggested that the ISF near capillaries would flow into both arterial and venous adventitial pathways after filtrated from capillary walls of the systemic vasculature.

To explore how the adventitial ISF along the pulmonary arteries entered the adventitial pathways along the pulmonary veins, we studied the temporal distribution characteristics of the adventitial FluoNa in the middle lobe of right lung where a pair of adjacent artery and vein could be found. At 50sec after the adventitial infusion of the FluoNa into the adventitial pathways of the saphenous vessels, the adventitial pathway of the pulmonary artery was stained but the adventitial pathways of the vein and the perivascular tissues were not (Fig. 6H, 6H1, 6K). This is significantly different from the results of hemoperfusion at 50sec after the intravascular FluoNa (Fig. S9A-C). Then, at 2min, the adventitial pathways of both the pulmonary artery and vein were stained (Fig. 6I, 6I1, 6K); at 3min, the adventitial pathways of the pulmonary artery and vein as well as the perivascular tissues in between have been stained together (Fig. 6J, 6J1, 6K). These results showed that the adventitial ISF along the pulmonary artery would enter the adventitial pathways along the pulmonary vein when an artery meets an adjacent vein.

### Effects of heart-targeted drug delivery for an adventitial ISF flow pathway

The potential function of the adventitial pathways as a novel route for drug delivery targeting the heart was explored. The effects on HR reduction in adventitial infusion group was found to be slower than that in intravenous injection group at 1-3min after the administration (Fig. 7A). The changes of Esmolol concentration detected in the myocardium of the ventricular apex coincided with the changes of the HR reduction by either adventitial infusion or intravenous injection (Fig. 7B). The Esmolol concentration in myocardium of intravenous injection increased rapidly and reached the peak at 1min after administration. The Esmolol concentration in myocardium by adventitial infusion reached the peak at 3min and remained higher that that by intravenous injection starting from 3min after administration.

**Figure 7.**
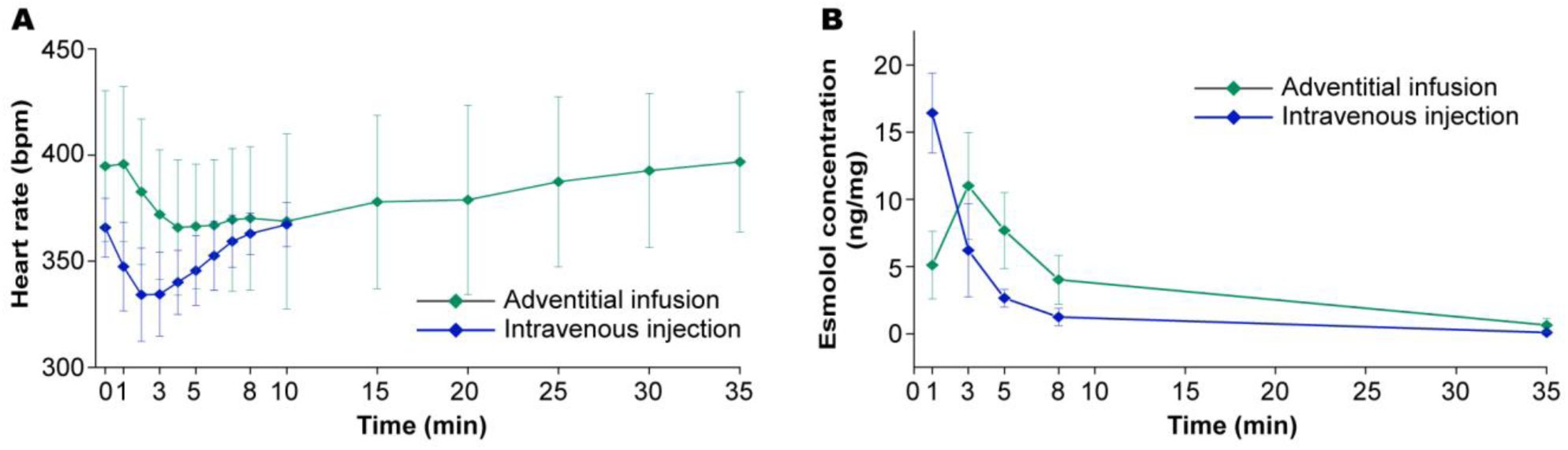
Reduction of the HR with Esmolol by adventitial infusion and intravenous injection. **A** The adventitial Esmolol caused a slow HR reduction at 1min after the infusion. The maximum reduction was at 4min, and the HR began to recover after 8-10min. In contrast, the intravenous Esmolol induced a rapid HR reduction immediately after the injection. The maximum reduction was at 2min, and the HR began to recover at 3min. **B** Detected by LC-MS/MS, the Esmolol concentration of intravenous injection increased rapidly and reached the peak at 1min. The Esmolol concentration of adventitial infusion increased slowly and reached the peak at 3min.

## Discussion

Building on previous findings on the long-range adventitial ISF flow along the larger-caliber vessels and its connections with the heart and lungs ^18–21^, our current experiments further identified that the ISF flows in adventitial matrix surrounded by layers of fascia (Figs. 2M, 2M1, 2O, 2O1, 2Q, S2A-C, S3B-D) and centripetally along the major arteries and veins of the systemic vasculature to the epicardium and myocardium (Figs. S1, S7), then into the lungs via the pulmonary arteries and returns the heart via the pulmonary veins (Fig. S8), which is neither perivascular tissues nor blood or lymphatic vessels. The regulatory mechanism of such adventitial ISF flow was found to be regulated by the heart and lungs and elucidated in the physiological or non-physical conditions of the rat model. Under physiological conditions of rats, the measured velocity of the speckle-like adventitial ISF flow along the femoral veins was positively correlated with heart rate, increased when holding breath, and became pulsative during heavy breathing. During respiratory cycle, each expansion of the lungs during inflation generates a centrifugal flow of adventitial ISF, while each contraction of the lungs during deflation generates a centripetal adventitial ISF flow. During cardiac cycle, each dilation of atria or ventricles generates a centripetal flow of adventitial ISF, while each contraction of atria or ventricles generates a centrifugal adventitial ISF flow. By repeated compressions of atrium or ventricles of the heart without respiration, the adventitial ISF was driven to flow centripetally along the femoral vessels (Fig. 5A-E1). By repeated ventilations of lungs without heartbeat, the adventitial ISF was driven to flow centrifugally along the femoral vessels (Fig. 5F-J1). It seemed that a respiratory movement of the lungs is able to retard the centripetal flow of adventitial ISF generated by the beating heart. As to the exchange with blood circulation, the ISF was found to flow longitudinally in the adventitial pathways along the vascular vessels, diffuse into the perivascular connective tissues, and return to blood circulation through the adjacent blood capillaries and lymphatic vessels during the long-range transport processes of either rats or rabbits ^9,21,35^. The detailed mechanism to maintain mass fluid balance among such an ISF transport, spread and redistribution system, blood circulation and lymphatic system requires more research. Here, our findings propose that the heart and lungs work together to drive the adventitial ISF to flow centripetally along the major systemic arteries and veins, and circulate around the pulmonary vasculature, comprising an adventitial ISF circulatory network, abbreviated as adventitial ISF circulation. Further identification of adventitial pathways along the arterioles, venules and capillaries in the lungs and other organs or tissues will provide the missing link to adventitial ISF cycle.

### The adventitial pathways were neither blood and lymphatic vessels nor the passageways in perivascular spaces

The adventitial pathways are the outermost connective tissues covering of vessels and neither blood and lymphatic vessels nor the passageways perivascular spaces in perivascular spaces ^18,21^. Even if the perivascular tissues were stripped, the ISF was still found to flow longitudinally along the vessels in the remaining adventitial tissues by the fluorescent tracer (Figs. S2B, S2B1, S3A-D1). When the remaining adventitial tissues were disrupted by collagenase I, the longitudinal flow of the fluorescent adventitial ISF along the vessels would be interrupted while the intravascular lumen was unobstructed (Figs. 2B-D, 2E, 2M, 2M1, 2O, 2O1, S2B-C1). Under the real-time stereofluorescence microscopy, the movement of the FluoNa in the longitudinal fluorescent lines along the side of the venous vessel clearly illustrated the continuous ISF flow in the remaining adventitial tissues (Fig. 2B, 2C, Video 5). As analyzed by real-time frozen sections, the adventitial pathways for the ISF flow, that were stained by the flow of the FluoNa from the distal (Figs. S2B, S2B1, S3A-D1), consisted of the outermost layer of fascia and the inner adventitial matrix, covering the media of either arterial or venous vessels. Unlike the conduit-like passageways in perivascular spaces next to the small intracranial vessels ^36^, the intrinsic structures of the adventitial pathways were composed of adventitial matrix, within which, the micro-CT found that there were approximate 2300-2800 fibers inside the femoral arterial or venous adventitial pathways, respectively (Fig. 2Q). However, neither fibers nor gel-like substances within adventitial matrix can flow freely along the vessels over a long distance. Hence, the boundary structures for adventitial ISF flow were the outermost fascia and innermost tunica media of blood vessels. We named this adventitial ISF flow pathway “interstitial matrix-membranes channel” ^9,20,21,37^.

The observed flow of adventitial ISF at different velocities suggested complex structures in the adventitial pathways as well. Within 20 seconds after adventitial infusion, fluorescein began to enter the adventitial channels, and the observed early rapid flow of the FluoNa represented a fast internal channel in the adventitial pathways. The subsequent speckle-like flow represented a slow internal channel in the adventitial pathways. Nevertheless, our current imaging techniques cannot further differentiate between the multiple fluorescein flow in these different internal channels. It is needed to develop a dynamic high-speed fluorescence imaging technology with higher resolution to visualize such diverse ISF flow in an adventitial pathway with multiple internal channels.

### The regulatory mechanism of adventitial ISF flow by the heartbeat and respiration

The dynamic patterns of the heart and lungs to regulate the adventitial ISF flow are unique. By measuring the motions of fluorescent speckles along the femoral veins under physiological conditions (Video 5), we found that the flow rate of adventitial ISF was positively correlated with the HR (Fig. 3B). The flow velocity of adventitial ISF increased during apnea and fluctuated by heavy breaths, and its fluctuation frequency was consistent with the breath rate (Fig. 3C, 3C1). By ventilation, it was showed that a large tidal volume can cause a pulsed centripetal flow of adventitial ISF in alive rats (Fig. 3C1, 3F1), while a pulsed centrifugal flow in dead rats (Fig. 3G1). By adjusting different parameters of mechanical ventilation in alive or dead rats, it was displayed that the frequency of the pulsed flow rate of the adventitial ISF was consistent with the respiratory rate (Fig. 3D, 3D1, 3E, 3E1); the inspiration and expiration (I/E) ratio (1:1, 2:1, 1:2) determined the descending and ascending branches of the velocity pulse (Fig. 3H, 3H1, 3I, 3I1). These findings showed the correlations between the changes in the adventitial ISF velocity and the heartbeat and respiration, clearly demonstrating that the adventitial ISF flow along the femoral vein was a synergistic result of the heart and lungs.

In our experimental observations, when the tidal volume of ventilation was less than 3 mL, almost no effect of breathing on the adventitial ISF flow rate can be found. When the tidal volume was greater than 4 mL, it can be displayed that the flow rate fluctuated with respiration. The proposed empirical dynamic equation (3) further illustrated the correlations of the adventitial ISF flow rate in an adventitial pathway with the cyclical motions generated by the regular heartbeats and irregular respirations (Fig. 3J-L, Video 6-8, Supplementary Note 1). In the present study, the continuous *T* represented the HIT of adventitial pathways along the femoral veins, while the fluid transportability of diverse adventitial pathways in other parts of various animals and humans requires further explorations.

The continuous cyclical diastolic and contractive motions of the heart and lungs were responsible for the to-and-fro flow of adventitial ISF. So how does each heartbeat or breathing motion of the lungs affect the adventitial ISF flow?

### The proposed regulatory mechanisms of the heart and lungs driving the adventitial ISF flow

To clarify the effects of each heartbeat or breath on the flow of adventitial ISF during cardiac or respiratory cycle, we performed experiments in dead rats by means of chest compressions, open-chest atrial or ventricular compressions, and mechanical ventilation, respectively. It was found that the speckle-like adventitial ISF along the femoral vein was “pulled” toward the heart when the atria (Fig. 4B2) or ventricles (Fig. 4C2) diluted, or the lungs (Fig. 4D2) contracted, whereas the adventitial ISF was “pushed” away from the heart when the atria (Fig. 4B1) or the ventricles (Fig. 4C1) contracted, or the lungs (Fig. 4D1) expanded. In freshly dead bodies of rats, repeated mechanical cardiac compressions can drive the adventitial ISF to move back and forth, and overall flow toward the heart (Figs. 4A, 5C). In contrast, repeated mechanical ventilations drove the adventitial ISF to move back and forth, and overall flow centrifugally along the femoral vein (Fig. 5H, Video 8). Due to the lack of quantitative observation methods of fluid flow in an arterial adventitial pathway, future studies are needed to reveal the effects of each stroke of the heart and lungs on the adventitial ISF flow along the systemic arteries as well as the pulmonary arteries and veins.

The mechanisms that every cyclical deformation of either the heart or lungs generates a pair of to-and-fro driving forces to “pull” or “push” the ISF to flow in the adventitial pathways along the systemic and pulmonary vascular tree are fascinating. Is it the perivascular pump driving the ISF to flow in the PVS of brain? There are two hypothesized causes of the perivascular pump, one is the shear force generated by the pulsating blood flow on the vessel wall, and the other is the mechanical pulsation generated after the heartbeat is transmitted through the rigidly connected vessel walls ^36,38^. To investigate the effects on the adventitial ISF flow by the pulsating blood flow and the mechanical pulsations of the vessel walls, we constructed an animal model with open ventricular chambers in air and meanwhile, the mechanical motions of the femoral vessel walls were detected by a laser vibrometer. Whether by repeated cardiac compressions or mechanical ventilations of lungs, no blood flow from the apex was observed during the experiments, while the measured arterial and venous pressure were almost close to zero and significantly lower than the physiological arterial and venous pressures (Fig. S6, Table S3). Under repeated cardiac compressions, the distal FluoNa was able to flow centripetally along the left femoral vessels (Fig. 5B, 5B1, 5E, 5E1). In contrast, the proximal FluoNa was driven to flow centrifugally along the left femoral vessels by repeated ventilation of the lungs (Fig. 5G, 5G1, 5J, 5J1). Moreover, we also found that the mechanical fluctuations of the vessel walls did not determine whether the FluoNa in the adventitial pathways flowed centripetally or centrifugally (Fig. 5A, 5D, 5F, 5I). Hence, these data showed that the driving forces for the continuous adventitial ISF flow come mainly from the expansions and contractions of the heart or lungs, rather than from the continuous pulsating blood flow or mechanical motions of the vessel walls. Of course, the other physiological factors that might involve in the adventitial ISF flow, such as the shear stress of blood flow, cardiac contractile strength, muscle contraction along the transport pathways, etc., need to be studied in the subsequent research.

Based on our previous and current findings ^18,20–22,39^, a working hypothesis was proposed to understand the mechanisms of the expansions and contractions of the heart and lungs “pull” or “pushing” the adventitial ISF flow centripetally or centrifugally along the femoral vessels: 1, When the heart and lungs expand or contract, the interstitial matrix within the heart and lungs will deform cyclically and work as a driver, named as “matrix or gel pump”. 2, The deformations of either cardiac or pulmonary matrix pump can generate at least two driving forces, the pressure gradients, and the fluctuating interfacial interaction forces of fluid in the diverse interspaces inside matrix ^21^. 3, The positive or negative pressure gradients generated by the atrial or ventricular matrix contracted, or the pulmonary matrix expanded will drive the adventitial ISF to flow along the vascular tree away from the driver or toward the driver. 4, Mediated by the intact adventitial tissues, the fluctuating interfacial interaction forces generated by the atrial or ventricular matrix diluted, or the pulmonary matrix contracted will “pull” the adventitial ISF to flow along the vascular tree toward the driver ^20,21^. In the human cadaver experiments, the “cardiac matrix pump” that was repeatedly compressed by an automatic cardiac compressor, had “pulled” the peripheral ISF in the thumb via the adventitial pathways into the epicardium^20^. In alive rabbits, the “cardiac matrix pump” had “pulled” the peripheral ISF via the adventitial pathways into the heart and entered the pericardial cavity, causing pericardial effusion^18,21^. As future works, the driving mechanisms of cardiac and pulmonary matrix pumps and the kinematics of adventitial pathways require more explorations.

As for the functions delivering fluid and dissolved constituents, we found that the adventitial infusion is significantly different from intravenous injection for drug delivery. The effects on the HR reduction of Esmolol by adventitial infusion was a slow and longer-drug-effect pattern compared to the rapid reduction by the intravenous injection (Fig. 7A). It was also found that the concentration of adventitial Esmolol was higher than that of intravenous Esmolol at 3, 5, 8 minutes, respectively (Fig. 7B). If Esmolol had been inside blood vessels, Esmolol would have been rapidly metabolized to inactive products by hydrolysis of the intraerythrocytic esterases in red blood cells^40^. The detected higher concentration of Esmolol in the ventricular tissues indicated that the adventitial infused Esmolol probably entered the myocardium through a fluid transport pathway that does not contain red blood cells. The presented pilot study may inspire the heart-targeted extravascular route of drug delivery in the future, especially for the drugs that seek to avoid rapid metabolism by erythrocytes.

In summary, our current findings showed that the heart and lungs play a synergistical role in regulating the ISF flow along the adventitia of vasculature throughout the body. The relationship of adventitial ISF flow with those around capillaries, and their mechanisms to maintain the homeostasis of internal environment between cells of diverse tissues or organs are particularly intriguing. The proposed matrix pump might inspire further research on the driving mechanisms of how the heart and lungs generate the forces of “pull” or “push” during a cardiac or pulmonary cycle. Under the actions of heartbeat and respiration, the centripetal to-and-fro flow of adventitial ISF along the systemic blood vessels may simultaneously participate in the drainage and perfusion of the ISF into and out of distal organs or tissues. For pulmonary circulation, the influence of the circulatory flow of adventitial ISF on the physiological and pathophysiological functions of the heart and lungs is a new research topic. Elucidating the relationship between adventitial ISF flow and vascular physiological functions and diseases will be an important advance in vascular biology. A specialized imaging technique is needed to study adventitial ISF flow along arteries. For drug delivery targeting the heart and lungs, the studies on the unique drug metabolism characteristics of adventitial ISF circulatory pathways may facilitate a novel therapeutic strategy for extravascular drug delivery and bring innovations in the fields of medical applications.

### Limitations of the study

1. Due to insufficient understanding of interstitial fluid flow along the systemic and pulmonary vasculature, this article can only focus on the regulatory mechanism of adventitial interstitial fluid flow by the heart and lungs, the directions of adventitial interstitial fluid flow and its potential drug delivery function. These insights will inspire future studies of physiological and pathological functions on the adventitial interstitial fluid circulatory network.
2. Although the graphic abstract looks “descriptive”, together with the empirical equation, it did reveal the regulatory mechanisms of adventitial interstitial fluid flow by the heart and lungs. The revealed regulatory functions will inspire future studies on the mechanical mechanisms of the heart and lungs on driving adventitial fluid flow.
3. Interstitial fluid flow in the adventitial matrix is passive and driven dynamically by heartbeat and respiratory movements. Limited by current imaging methods, we can only observe the patterns of fluid flow in venous adventitial pathways. The imaging methods on observing the dynamic flow of adventitial fluid through arterial adventitia needs further studies.
4. Conventionally, the adventitial matrix is a gel substance, through which, fluid cannot flow freely over long distances. Our current data verified that it is the very adventitial matrix, and its surrounding fascia are involved in fluid transport. Together with our previous studies, it is reasonable to speculate that there must be a space for fluid flow in the adventitial matrix paved by layers of fascia. However, clear imaging of fluid flow in different spaces within the adventitial pathways requires more research, especially high-resolution imaging techniques when fluid flows at high speeds.

## Author contributions

HY.L. conceived and developed the original ideas, concepts and theory of the interstitial fluid circulatory network. HY.L., H.L., C.M., and FS.J. designed the experiments. HY.L. wrote the paper. B.L., WQ.L., X.Q., HY.L., JZ.L., Z.H., X.Y., DH.H. performed the experiments. Y.H., B.L., H.L. calculated the velocity. HY.L., H.L., Y.H., B.L., carried out the empirical formula. JP. Z, HY.L., B.L., TT.L., performed the MRI experiments. B.L., CZ.Y., ZM.L., and T.G. performed the spectral micro-CT experiments. HY.L., C.M., FS.J., J.H., and ZJ.Z. analyzed the imaging data. WenQing.L., HY.L., F.W., and L.L analyzed the data for drug delivery experiments. HY.L. and B.L. prepared pictures and videos.

## Acknowledgements

We thank Professors Guanhua Xu, Yixin Zeng and Kaixian Chen, Weiwu Hu, Bo Song, Saicang Lobsang Hwardan Chegynima, Jamyang Danchour, Angwen zhabar, Andreas Sammer and Georg Feigl for their fruitful discussions. We thank Weihua Wang, Fang Wei, Yu Tang in center of pharmaceutical technology of Tsinghua University for their help in LC-MS/MS.

## Methods and materials

### Subjects and the selection of the imaging tracers

All animal experiments were approved by the Institutional Animal Care and Use Committee of the Institute of Basic Medical Sciences Chinese Academy of Medical Sciences, Peking Union Medical College (No. ACUC-A02-2021-017) and performed according to institutional guidelines and research protocols approved by the Institutional Animal Care and Use Committee of the Institute of Basic Medical Sciences Chinese Academy of Medical Sciences, Peking Union Medical College. A total of 189 male Sprague‒Dawley rats, 300-350 g in weight (HFK Bio-Technology, Beijing, China), were used and housed in pathogen-free conditions with 12 hours of continuous light and 12 hours of continuous darkness at the Laboratory Animal Center of the Institute of Basic Medical Sciences Chinese Academy of Medical Sciences. The procedures designed minimize animal suffering and respect the 3Rs principles. The rats were anaesthetized with isoflurane (1%) in 1 L/min oxygen or intraperitoneally anaesthetized (pentobarbital 50 mg/kg) during experiments. Body temperature was maintained at 37.5 °C with a rectal probe-controlled heated platform. A ventilator (RoVent Jr., Kent Scientific, USA; V100, YUYAN, China) was used to regulate the frequency, tidal volume, and I/E (inhalation/exhalation) of breath. The rats were euthanized by carbon dioxide according to the guidelines of the American Veterinary Medical Association (AVMA).

Using fluorescent stereomicroscopy (Axio Zoom.V16, Zeiss) and the spectral micro-computed tomography (SMCT) ^41–43^, we examined which tracers were able to flow in the adventitial pathways along the vessels of the lower limbs in rats, including several large (>1 kDa) and small (<1 kDa) molecular weight, water– and lipid-soluble tracers (Table S4). It was found that the fluorescein sodium (FluoNa) can be used for *in vivo* real-time fluorescent imaging with a higher spatial resolution than Rhodamine B and Indocyanine green, and the silver nitrate can be used for *in situ* imaging within tissues and organs (Fig. 1, Video 12). The gadolinium-diethylenetriamine pentaacetic acid (Gd-DTPA) was used in MRI experiments. The FITC-dextran (70,000 dalton) was used to visualize the lymphatic vessels accompanying the femoral vessels.

### Methods for adventitial infusion and intravenous injection by fluorescent, paramagnetic, or silver tracer

For intravenous injection, the selected tracer was injected into the saphenous vein at the level of right knee. For adventitial infusion by fluorescent tracer, 4µL solution of FluoNa (diluted to 0.1 g/L in normal saline (NS)) was dripped onto the adventitia on the saphenous or femoral arteries or veins of the lower limbs, axillary artery and vein of the upper limbs or the left common carotid artery in the middle neck or the anterior descending arteries near the apex of the heart using the tip of the 10 µL pipette. 40µL FluoNa (0.01 g/L in NS) was used for intravenous injection.

Consistent with the imaging methods in our previous studies ^18,21,44^, the adventitial infusion for MRI or SMCT was the hypodermic injection of the Gd-DTPA or silver tracer into the right or left ankle dermis. This method is equivalent to injecting the tracer into the perivascular tissues of the accompanying saphenous artery and vein under ankle dermis. The duration for the hypodermic injection into ankle dermis was 5-10sec. 200µL Gd-DTPA (0.5 mmol/mL in NS, MedChem Express, Quality Research, Zhuozhou, Hebei, China) was used for adventitial infusion or intravenous injection, respectively. 400µL silver nitrate solution (10% in deionized water) was injected into ankle dermis for the cardiac perfusion by adventitial infusion and 1mL silver nitrate (2% in deionized water) for intravenous injection. 4µL silver nitrate solution (10% in deionized water) was dripped onto the adventitia on the saphenous arteries and veins in the lower limbs to detect whereabouts the adventitial ISF flowed along the saphenous vessels. The angiography was performed by injecting the tracer into the tail vein.

### Image acquisition by MRI

Imaging was performed by 9.4T MRI scanner (Biospec 94/30 USR Bruker, Ettlingen, Germany) with a rat body volume coil. A respiratory sensor (SA Instruments, Stony Brook, NY, USA) was placed under the abdomen to monitor the respiration rate. Scanning parameters were adjusted to obtain a high spatial resolution. The coronal images were collected with a 3D FLASH sequence: TR=11ms, TE=1.84ms, flip angle=10°, FOV=90×31×23mm, matrix size=510×176×131, resolution=176μm isotropic, number of averages=1, acquisition time was 5 min. Immediately after the administration of the Gd-DTPA, each rat was dynamically scanned. The raw data were analyzed at a Dell Precision Tower workstation (T7910) with multiplanar reconstruction (MPR) and maximum-intensity projection (MIP) reconstructions. 5 rats were used for adventitial infusion and 3 rats for intravenous injection.

### In vivo fluorescence imaging of the adventitial pathways in the lower limb

After the skin along the vessels of the limbs opened surgically in 6 rats, the FluoNa was administrated by adventitial infusion on the saphenous vessels or intravenous injection into the saphenous vein, respectively. The real-time flow of the fluorescent adventitial ISF along the arterial and venous vessels in the limbs (Fig. 1B, 1G) was recorded by the fluorescence stereomicroscope with a high-sensitivity camera (Prime BSI Scientific CMOS, Teledyne, USA). After recording, sections of the stained arteries and veins at the distal and proximal injection site (Fig. 1C, 1D, 1H, 1I) were sampled for the real-time frozen fluorescence analysis.

In another 5 rats, a 3-5mm segment of the right or left femoral vessels was isolated and exposed to air by surgically stripping their perivascular tissues (Fig. 2A, 2B, 2C, S3A, S3A1) while the intravascular blood flow was kept intact. The continuous flow of the adventitial FluoNa along the isolated femoral artery and vein was clearly recorded when the FluoNa was administrated onto the adventitia of the distal saphenous vessels. After recording, the distal femoral vessels stripped of perivascular tissues (Fig. 2M, 2M1, 2O, 2O1, S3B-D1) and the proximal femoral vessels with perivascular tissues (Fig. 2H, 2H1, 2J, 2J1, S2B, S2B1) of the same rat were sampled for the real-time frozen fluorescence analysis, respectively.

After the adventitia along the isolated femoral vein and artery was bathed by 10µL solution of type I collagenase (0.1 g/mL in NS, Sigma-Aldrich) for 30min in the other 6 rats, it was observed whether the distal FluoNa could pass through these disrupted adventitial pathways (Fig. 2D, S2C, S2C1). By intravenous injection of the FluoNa into the lumen of the distal saphenous vein (2/6 rats) or the contralateral saphenous vein (2/6 rats), it was observed whether the blood flow of these femoral vessels was unobstructed (Fig. 2E). In the last 2/6 rats with the disrupted adventitia, the femoral vessels were sampled for the frozen fluorescence analysis after the adventitial infusion of the FluoNa (Fig. S2C, S2C1).

In the control group of 3 rats by adventitia-infused NS, the corresponding tissues in the limbs, abdomen and thorax were sampled for frozen fluorescence or histological analysis. To visualize the lymphatics, 200µL solution of FITC-dextran (1%, 70kDa, Sigma-Aldrich) was injected hypodermically into the ankle dermis (Fig. S3K-K2).

### Methods for measuring the flow rate of early rapid flow and subsequent speckle-like flow by fluorescence imaging methods

Limited by the spatial and temporal resolution of MRI, we investigated the flow rate of adventitial ISF using fluorescein. Immediately after the FluoNa administration by real-time fluorescence stereomicroscope with high-sensitivity camera, the early and subsequent flow of the fluorescent adventitial ISF was recorded dynamically along the femoral vessels with stripped perivascular tissues. The distance between the administration site and the observation point and the early time of fluorescein appearance on the femoral vessels were recorded in 3 rats. The speckle-like flow along the femoral veins was recorded in a total of 24 rats (Table S2).

We developed a method of Speckle tracking velocimetry (STV) to measure the continuous FluoNa flow by speckle pattern translation recording, which has been extensively used in measuring displacements and velocities ^25,27,28^. The STV consists mainly of two steps: 1) Pre-processing; 2) Velocimetry. 1) By analyzing the data of the fluorescent adventitial ISF flow, we found several difficulties in measuring velocities in the videos, such as the low-frequency background light flickering and the quiver of the vessel itself. Therefore, in the first step, we designed an algorithm to overcome these difficulties based on some image processing methods, such as high-pass filter, image registration, and mean-image subtraction. 2) For the velocimetry, the region of interest was divided into small patches (as the rectangle in Fig. S5A). Afterwards, for each small patch, cross-correlation ^26^ was performed to estimate the most probable displacement within its surrounding area in the next frame. Then, displacement per time between adjacent frames yielded the velocity for each patch (as the vector field in Fig. S5A). The velocity field of the continuous adventitial ISF flow along the femoral vein was analyzed by the STV in 3 live rats (Fig. S5B, S5C). The real-time flow of adventitial ISF along the femoral veins was recorded by the fluorescence stereomicroscopy while recording synchronized measurements of the electrocardiogram (ECG) and the respiratory cycle by animal physiological monitoring device (PowerLab, AD Instruments, Australia). The speed change curves were found to match the changes of heartbeats and respiration.

The flow rate changes of the FluoNa along the femoral veins under physiological and non-physiological conditions were further investigated by STV. The changes of the adventitial ISF flow velocity at different heart rate (HR) were recorded in a typical rat (Fig. 3B). The HR was lowered by tail vein injection of Esmolol (2.0-3.0 mg/kg/min). The effects of the different respiratory parameters on the adventitial ISF flow were investigated in 6 live (Fig. 3D1, 3F1, 3H1) and 6 freshly dead rats (Fig. 3E1, 3G1, 3I1), such as respiratory rate (RR), inspiratory expiratory ratio (I/E) and tidal volume (TV). The freshly dead rats referred to the rats that were euthanized by carbon dioxide and used for the experiments within 1 hour after the heartbeat and breathing completely stopped for 15 minutes (detected by ECG and respiration signal monitoring of PowerLab). Two pressure-measuring guidewires were placed in the contralateral iliac vein and artery of each rat. The systolic and diastolic arterial pressures, venous pressures of iliac vessels were measured under different respiratory parameters (RR: 30-60-90bpm, I/E: 1:1-1:2-2:1, TV: 4.0-6.0-8.0mL), respectively. The changes of the arterial and venous pressures were recorded by PowerLab. The changes of breathing (Resp.) were the measured pressure on the surface of the body of the rats by the breathing band sensor.

### Qualitative observation on the circulatory pathways of adventitial ISF flow along the systemic and pulmonary vasculature by the fluorescence imaging

To depict the circulatory pathways of adventitial ISF along the systemic vasculature, we tracked the longitudinal flow of adventitial ISF along several major systemic arteries and veins by the adventitial infusion of the FluoNa onto the right saphenous vessels, axillary vessels, and the left carotid artery of the neck in 9 rats, respectively (Fig. S7A). The distal and proximal ends of the vessels around the injection site were sampled for the real-time fluorescence or histological analysis. The fluorescently stained right anterior vena cava (AVC), inferior vena cava (IVC), and aorta were sampled and compared with those of adventitial infusion by NS.

To delineate the cycle of adventitial ISF flow along the pulmonary vasculature, the pulmonary artery, vein, and the surrounding tissues between the artery and vein at the level of the right middle lobe of lungs were sampled at 50sec, 2min, 3min after the adventitial infusion of the FluoNa into the right ankle dermis in 9 rats, respectively. In other 4 rats, the FluoNa was administrated on the surface of the root of the pulmonary artery or a segment of the pulmonary vein in the middle lobe of right lung, respectively. At 10sec after the administration, the movement of the FluoNa was recorded by stereo fluorescence microscope.

To disclose the flow direction of adventitial ISF along the coronary arteries of a beating heart, the FluoNa was dropped into the superficial tissues on the ventricular apex of 3 rats, and dynamically recorded with a high-speed camera.

### Ex vivo fluorescence imaging and histological analysis

Frozen fluorescent section slices (4μm thickness) were obtained using calibrated vibratome (VT1200S, Leica). One section was selected every 5 sections for each sample of the arterial and venous vessels, heart, or lungs. All slices were imaged by the fluorescence microscope (Scope.A1, Zeiss, German) with a digital camera (Axiocam 506, Zeiss, German). Exposure time, magnification, and luminous intensity under a dark environment were kept the same for all groups. Quantification of fluorescence intensity of the slices was performed by ImageJ (1.8.0). In the cross-sectional images of the vascular vessels of each rat, the arterial, venous walls, and perivascular tissues were equally divided into 5 regions. The maximum fluorescence intensity value in each region was recorded. There were 3 rats in each experimental or control group. Thus, a total of 15 fluorescence intensity value of the arterial, venous walls, and the surrounding tissues were obtained for each group. The same thresholds were used for the slices of each group. The frozen slices were also studied by Elastica van Gieson staining. The hematoxylin and eosin (H&E) staining and the immunostaining of the antibodies against CD31 (Servicebio, Wuhan, China) were performed according to routine procedures.

### Methods for investigating the effects of each heartbeat or breath on the adventitial flow in freshly dead rats

Under physiological conditions, it was found that the adventitial ISF flow was continuous centripetally along the femoral vessels and fluctuated with respiration motions at high tidal volumes. To investigate the effects of each heartbeat or breath on the adventitial ISF flow, we performed the following experiments by means of chest compressions, open-chest atrial or ventricular compressions, and mechanical ventilation in freshly dead rats, respectively. The flow rate of the adventitial ISF flow along the femoral veins was measured by the STV as well. The systolic and diastolic arterial pressures, venous pressures of iliac vessels were measured under the respiratory parameters (RR: 90bpm, I/E: 1:1, TV: 6.0mL). When cardiac and respiratory arrest, all dead rats were given heparin (125 IU/kg) for anticoagulation via the tail vein before the following experiments.

By chest compressions at a frequency of about 30 per minute, the flow rate of adventitial ISF was recorded in 3 freshly dead rats (Fig. 4A). The duration of one-shot compression was around 1sec. By surgically opening the chest cavity, the effects of each contraction and dilation of atria or ventricles on the adventitial ISF flow were observed when the atria or ventricles were directly compressed in 6 rats, respectively (Fig. 4B, 4C). The compressions of the atria or ventricle were performed by a digital force gauge (HF-10, HighTec, China) at the force around 0.5 N. The duration of one-shot compression was around 10sec. The free relaxation time of the atrium or ventricle after one-shot compression were about 10-15 sec. The ECG signals were detected when the heart, atria or ventricles of dead rats were compressed, representing the compressions.

In another 3 freshly dead rats, the effects of each inflation and deflation of the lungs on the adventitial ISF flow were recorded when the lungs were ventilated by a 10mL syringe (Fig. 4D). The inflation and deflation volumes were both 6mL. The duration of inflation time was around 10 sec, and the deflation time was about 10-15 sec.

### Methods for investigating the effects of repeated cardiac compressions or ventilations on the adventitial flow in freshly dead rats with opened ventricular chambers

The effects on the adventitial ISF flow by the pulsating blood flow and the mechanical pulsations of the vessel walls were studied in 16 freshly dead rats who were given heparin (125 IU/kg) for anticoagulation via the tail vein as well. After preparation, the apex of the heart was surgically removed in all dead rats to expose the left and right ventricular chambers to the air, and the heart was repeatedly compressed to drain the residual blood in the vascular vessels until no more blood was flowed out. Subsequently, the mechanical motions of the femoral vessel walls were detected by a laser vibrometer (OFV-5000, Polytec). The systolic and diastolic arterial pressures, venous pressures were also measured under certain respiratory parameters (RR 90bpm, I/E 1:1, TV 6.0mL). Two pressure-measuring guidewires were inserted from the open left ventricular chamber into the abdominal aorta and from the open right ventricular chamber into the abdominal inferior vena cava, respectively. The changes of the arterial and venous pressures were recorded by PowerLab. The adventitia of the right femoral vessels was disrupted by collagenase I (10μl, 0.1 g/mL in NS) and the left femoral vessels were bathed in NS solution. The flow of the adventitial ISF along the right or left saphenous or femoral veins was recorded dynamically by fluorescence stereomicroscopy and the velocity was measured by STV.

After administration of the FluoNa at the distal end of the adventitial pathways along the left and right saphenous vessels, the heart was pressed at 30-60bpm for 30 minutes in 6 rats while the lungs were not ventilated (Fig. 5C). After administration of the FluoNa at the proximal end of the adventitial pathways along the femoral vessels, the lungs were ventilated for 30 minutes in another 6 rats while the heart was not compressed (Fig. 5H). The results of the adventitial ISF flow along the right and left femoral and saphenous vessels were recorded by fluorescence stereomicroscopy, respectively.

To clarify whether the forces generated by the repeatedly ventilated lungs drive the adventitial ISF flow through the conduction of the pulmonary artery or veins, we continued to use vascular clamps to clamp the root of the main pulmonary artery or the thoracic segment of the inferior vena cava in the last 4 rats, respectively.

### Detection of the distributions of silver nitrate in tissues by SMCT

To determine whereabouts the adventitial ISF flowed along the saphenous vessels or entered the heart, we investigated the adventitial pathways in the lower limb and the cardiac perfusion by adventitial infusion or intravenous injection of the silver nitrate, respectively. 3 rats were used to detect the locations of the adventitial infusion of 4µL silver nitrate solution that was dripped onto the adventitia on the saphenous arteries and veins. The cardiac perfusion by adventitial infusion of silver nitrate was used in 5 rats and the intravenous injection in another 5 rats. If the rats died during the injection, the repeated chest compressions (400 bpm) were given immediately to ensure the circulate of the silver nitrate in the blood to reach 3 minutes. The rats were sacrificed at 3min after the administration. The samples of the proximal femoral artery and vein (2cm from the infusion site), the heart and lungs were sampled. All samples were treated with alcohol gradient dehydration (30%, 50%, 75%, 90%, 100%) before SMCT scanning.

The samples were imaged using a photon counting detector (PCD) based spectral micro-CT scanner (The Institute of High Energy Physics, Chinese Academy of Sciences, China). The PCD (XCounter, Sweden) uses a CdTe crystal with a thickness of 0.75 mm as the sensor material. The detector contains 2,048×512 pixels with a pixel size of 100 µm. It has two energy thresholds and can detect photons with energy range from 10 keV to 160 keV. By setting the thresholds, two energy window data can be obtained in one CT scan. ^41–43^.

### Detection of the intrinsic structures in the adventitial pathways along the femoral artery and vein by micro-CT

The femoral vessels of 3 rats were sampled and treated with alcohol gradient dehydration (30%, 50%, 75%, 90%, 100%) before micro-CT scanning. High-resolution 3D X-ray micro-CT (nanoVoxel-3000 series; Sanying Precision Instruments Co., Ltd.) was used to get 1800 pics grayscale images of each sample ^20,45^. The images were captured on a detector 2940×2304 pixels, with an exposure time of 0.45sec, a voltage of 60.0 kV, a current of 50 μA and a voxel size of 0.85μm. The software VOXEL RECON (Sanying Precision Instruments Co., Ltd.) was adopted to reconstruct algorithm and correct image data.

The numbers of the adventitial fibers between the fascia and the tunica media of the femoral arteries and veins were estimated in the reconstructed 3D images of micro-CT. The area of the adventitial fibers and its surrounding fascia along the femoral artery and vein were identified and extracted. The minimum intensity value of the fibers between the fascia and tunica media was set for connected component analysis. The diameter of the fibers was set to give the cross-sectional area of a fiber. The area of the connected components was then divided by the area of the fibers to obtain the numbers of fibers in the connected components. Then, the numbers of bundles in all slices were counted. The maximum, median and minimum values of fibers in adventitial pathways along both femoral artery and vein were calculated respectively.

### Esmolol administration and concentration in myocardium analysis

The changes of the HR by adventitial infusion or intravenous injection of Esmolol was recorded in 10 alive rats, respectively. The dosage of Esmolol for adventitial infusion or intravenous injection was 0.5 mg/kg and adjusted according to each rat weight ^46^. After the adventitial infusion or intravenous injection of Esmolol, another 50 rats were sacrificed at 1, 3, 5, 8, 35min, respectively. The concentration of Esmolol in ventricular muscles was detected by LC-MS/MS. The samples were homogenized by the grinder (Tiangen-Osey50) with 1 µL/mg of methanol acetonitrile (50:50) % (v/v), centrifuged at 4000 rpm (Eppendorf 5810R) for 20 minutes, and the supernatant was taken for detection. Measurements of the Esmolol were accomplished on a Shimadzu LC-2-AD XR equipped with a binary pump, a degasser, an auto sampler/injector and a column heater (Shimadzu, Kyoto, Japan), coupled to an AB SCIEX QTrap 5500 Mass Spectrometer equipped with an ESI ionization source (AB SCIEX, USA).

### Statistical analysis

Statistical analyses were performed using Origin 2022 software. Paired samples t-test was used to compare the fluorescence intensity of two different points in a single group and two-tailed Student’s t-test was used between two groups. The exact P values were calculated at a 0.05 level of significance and stated in the figure legends.

